# Environmental effects on constructed wetland microbial diversity and function in the context of wastewater management

**DOI:** 10.1101/2025.01.14.633069

**Authors:** Sandrine Grandmont-Lemire, Bob Gearheart, Catalina Cuellar-Gempeler

**Affiliations:** Chilkat Indian Village; Arcata Marsh Research Institute (AMRI), Arcata, CA; Cal Poly Humboldt, Arcata, CA

## Abstract

Considering temporal and spatial change in biodiversity-ecosystem function (BEF) relationships is critical to predict and manage ecosystem services, especially in human mediated and impacted ecosystems. We propose that species responses to seasonal change and spatial distributions can act as a laboratory to reveal diversity-function relationships with management implications. This study investigates the relationship between bacterial diversity and ammonia removal function in a wastewater secondary treatment constructed wetland system. We took 8 samples across a system of 6 interconnected ponds, from August 2019 to February 2020, at the Arcata Wastewater Treatment Facility (AWTF), in Coastal Humboldt County (California, USA). We used 16S rRNA gene amplicon sequencing to measure bacterial diversity and composition, and an ammonia electrode probe to measure NH_4_ at the influx and efflux positions of each pond. We found a significant negative relationship between ASV richness and ammonia removal, suggesting that nitrifying and denitrifying bacteria are poor competitors, known a negative selection effect. Bacterial richness effect on ammonia removal was strongest, followed by direct effects of season on richness and location on function, based on structural equation modeling. We identify taxa associated with function that may influence management strategies, including Planktophila, Legionella, Sulfurimonadaceae and Sporichtyaceae that thrive in ponds located after chlorination steps. This result challenges the traditional wastewater treatment reactor paradigm to reveal negative BEF relationships that appear stronger than environmental influences. By expanding our views of BEF relationships, we can further unravel how community diversity and composition influence ecosystem processes in natural and humanized systems.

**IMPORTANCE:** This study sheds new light on how biodiversity impacts ecosystem functions in human-made environments, specifically wastewater treatment systems. By examining bacterial diversity and ammonia removal efficiency across interconnected ponds, we challenge the conventional assumption that more species always lead to better ecosystem performance. The surprising finding that higher bacterial diversity can reduce ammonia removal efficiency (due to competition among key bacteria) offers fresh insights into how microbial communities work. This understanding is critical for improving wastewater treatment and designing systems that maximize efficiency. Moreover, identifying specific bacteria linked to ammonia removal provides practical information for better managing and enhancing treatment processes. By broadening how we think about the relationship between biodiversity and ecosystem function, this study offers valuable tools for both scientists and environmental managers working to balance human impact with ecosystem health.

## INTRODUCTION

Biodiversity and ecosystem functioning (BEF) relationships have been an area of inquiry for community and ecosystem ecologists for the past three decades (Cardinale et al. 2012, Naeem et al. 2012, Isbell et al. 2017). The literature generally supports positive BEF relationships, with abundant evidence focusing on primary productivity or biomass accumulation as the central funciton (Balvanera et al. 2006, Cardinale et al. 2011, Isbell et al. 2017). The BEF field has now been extended to understanding the impact of climate change and human activities on biodiversity for terrestrial, marine, and aquatic habitats as well as serving as an urgent call for conservation to maintain ecosystem functions (Loreau and Hector 2001, Cardinale et al. 2012, Tilman et al. 2014). However, this positive BEF trend is not unique, with field studies demonstrating diverse trends in biodiversity and function relationships (Rychtecká et al. 2014, van der Plas 2019, Hagan et al. 2021). Importantly, we have a poor understanding of how non-positive BEF relationships respond to climate and human induced change, especially for ecosystem functions like degradation (due to negative selection effects, (Jiang et al. 2008, Peter et al. 2011, Cuellar-Gempeler et al. in review), pathogen defense (due to dilution effects, Schmidt and Ostfeld 2001, Hambäck et al. 2014) and grassland productivity (due to priority effects, Catano et al. 2023). Instead of searching solely for a generalized BEF relationship, we should unveil how different BEF relationships respond to environmental change and move towards a more dynamic view of biodiversity-function relationships (D’Andrea et al. 2024).

A dynamic view of diversity is well supported by our understanding of assembly processes such as abiotic filtering, species interactions, and habitat patch spatial arrangement is well documented, and yet the functional outcomes are less clear (Leibold et al. 2017, Bannar-Martin et al. 2018, Chase et al. 2020). Abiotic factors along a gradient of increasing stress can reduce diversity through habitat filtering, favoring species with tolerance traits that enable them to maintain viable populations (Kraft et al. 2015), while also selecting the functional traits of the community (de Bello et al. 2012, Zobel 2016). In turn, species interactions can exclude species and their function through mechanisms like competitive exclusion, but can also modify the functional output of coexisting species through co-regulation (Greiner La Peyre et al. 2001, Snel et al. 2004). When considering communities interconnected in space (metacommunities), it is key to consider how space can influence diversity via dispersal limitation and heterogeneity (Leibold et al. 2004, Hanson et al. 2012, Nemergut et al. 2013), yet the functional consequences of these spatial processes are seldom known. Overall, predicting the combined effects of these mechanisms on diversity and function is challenging, as both can be influenced by the same underlying variables (Krause et al. 2014). We can address this challenge by leveraging BEF relationships along environmental gradients to make quantitative predictions (Tilman et al. 2014, Wardle 2016, Eisenhauer et al. 2016, De Laender et al. 2016). Such approach would allow us to address a key remaining question: whether environmental conditions and assembly processes exert a stronger influence on function than diversity itself. This raises the possibility that environment has non-random effects on species contributions to function (Suding et al. 2008, Gonzalez et al. 2016) not through shifts on population density but via per capita functioning (De Laender et al. 2016).

To address this question, this study focuses on the bacterial communities within the Arcata Wastewater Treatment Facility (AWTF) which is composed of oxidation ponds and constructed treatment wetlands used to remove waste and toxins from wastewater before releasing it into the Humboldt Bay. The AWTF is composed of three phases of constructed ponds categorized as a free water hydraulic design (Supplementary Methods). The ponds are connected in series to facilitate wastewater processing, with particular focus on Ammonia removal. Ammonia is one of the principal pollutants found in wastewater, originating from industrial and house-cleaning chemicals, amino acid products, and urine in sewage (Minocha and Rao 1988) with consequences for algal blooms, effluent quality, and public health (Vitousek et al. 1997, Bodelier et al. 2013). While traditional wastewater systems rely on costly energy inputs to function (Knowles 1996, Lu et al. 2014), constructed wetlands provide a series of habitats where macrophyte and microbial communities perform metabolic processes in relatively natural conditions to remove ammonia from the untreated or previously treated wastewater (Dong and Reddy 2012, Liang et al. 2016, Doering et al. 2021, Sroufe and Watts 2022). In oxidation ponds, algae and macrophytes can uptake ammonia directly, while providing physical substrate, carbon sources and oxygen to support bacterial nitrification. Additionally, a key distinction between these systems is that traditional wastewater systems consist of a fixed number of reactors, which are relatively isolated from the environment, whereas constructed wetlands often feature spatially explicit interconnected ponds directly influenced by surrounding environmental conditions, making this an intriguing system to evaluate the effect of location and season on bacterial diversity and ammonia removal.

The metabolic pathways central to ammonia removal from wastewater are nitrification and denitrification, as well as uptake by algae, aquatic macrophytes and bacteria. Nitrifying bacteria metabolize the ammonia into bioavailable nitrates which photosynthetic organisms require for growth, including primary producers like plants and phytoplankton. While primary producers can utilize ammonia as a nutrient source, wastewater contains it in excess, and plants or phytoplankton instead require a balanced nitrate-to-ammonia ratio to optimize ammonia uptake efficiency (Errebhi and Wilcox 1990). Following nitrification, a large portion of the nitrate is reduced to molecular nitrogen and released into the atmosphere by denitrifying bacteria (Bodelier et al. 2013), thus completing ammonia removal from the system. Importantly, bacteria responsible for ammonia removal require large and complex enzymes, tend to grow slowly and are poor competitors, raising questions of their persistence in the system and maintenance of their critical ecosystem function in wastewater impacted habitats (Prosser 2007, Hoffman et al. 2023, Zhu et al. 2023).

Bacteria-driven ammonia removal may be influenced by seasonal environmental change and the spatial distribution of the wastewater treatment wetlands and we consider four hypotheses that can explain the emergence of negative (Hypotheses 1 and 2), positive (Hypothesis 3) and neutral (Hypothesis 4) BEF relationships. First, different habitat conditions across the constructed ponds can generate heterogeneity and dispersal limitation that may support ammonia removal via regional coexistence of otherwise poor competitors (Hypothesis 1). Second, we hypothesize that season could result in habitat filtering due to low temperatures in winter, benefiting cold tolerant taxa that may or may not be contributors to ammonia removal (Hypothesis 2). A third alternative would result in a positive BEF relationship where bacteria contributing to ammonia removal benefit from increased diversity that supports facilitative or mutualistic relationships between bacterial taxa, via cross-feeding (Smith et al. 2019, Giri et al. 2021) or complementarity (Loreau and Hector 2001, Tilman et al. 2014, Barry et al. 2019, Hypothesis 3). Lastly, taxa could be functionally redundant and achieve ammonia removal efficiently regardless of their diversity or composition (known as functional redundancy, Walker 1992, Lawton 1994, Loreau 2004, Hypothesis 4). Understanding the mechanisms that sustain taxa capable of ammonia removal will enable us to more accurately predict changes in diversity and function across environmental and biological gradients.

The central goal of this study is to investigate whether species responses to seasonal environmental change and habitat spatial distributions reveals diversity-function relationships with management implications. Specifically, we ask how location and season influence the relationship between bacterial communities and their function in the context of Ammonia removal within the AWTF. Based on the specific requirements of nitrification and denitrification processes, we expect to find a negative relationship between bacterial diversity and ammonia removal. We also hypothesize that bacterial diversity will have a stronger effect on function than environmental conditions and spatial relationships because of the strong metabolic component of this process. To understand the underlying processes, we identified bacterial taxa associated with ammonia removal and explored their potential functional and coexistence traits. We assess community level responses to season and location in aims to propose strategies that maximize ammonia removal from wastewater before the bay discharge point. By expanding our understanding of microbial BEF relationships in these wastewater treatment wetlands we can further unravel dynamics in natural and humanized systems with the goal of predicting, managing, and sustaining adequate ecosystem functions.

## METHODS

This study was conducted from October 2019 to February 2020 at the Arcata Marsh located in Coastal Humboldt County (California USA). These constructed wetlands are composed of ponds connected in series: Oxidation pond 1 and 2, parallel treatment wetlands, Allen Pond, Gearheart Pond and Hauser Pond (Fig 1, Supplementary Materials). There is a chlorination step before Oxidation pond 1, one after the parallel treatment wetlands and a last one before the Bay discharge point. Ponds vary from open water (Oxidation ponds) to semi-vegetated (Allen, Gearheart, Hauser ponds, Supplementary Materials).

**Figure 1.**
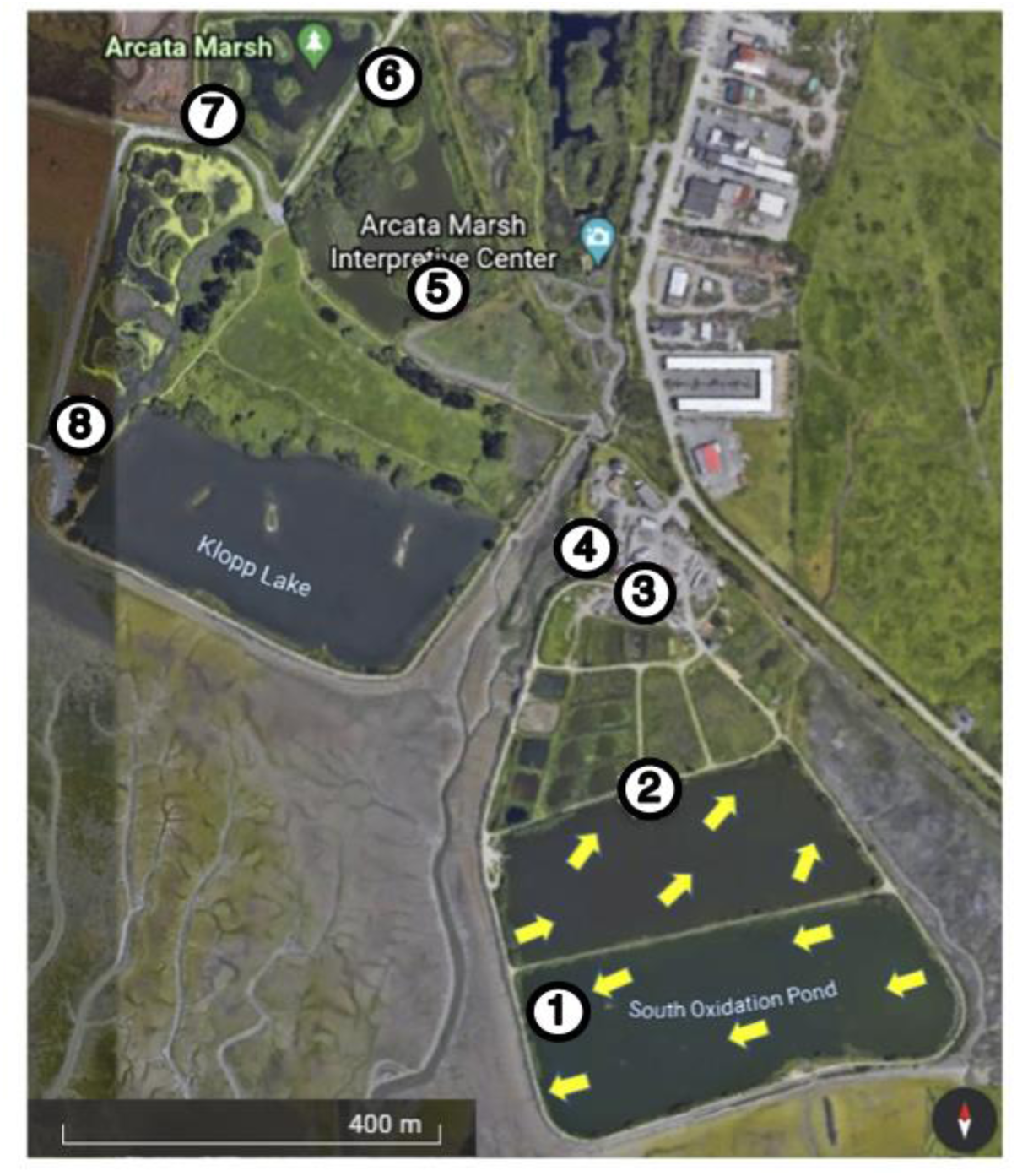
Map of the study site and sample locations. Numbers indicate sample locations at weirs that access the influent and effluent for each pond. For Oxidation Ponds (1, 2), yellow arrows represent sampling locations as distributed along the water flow from influent to effluent. The samples from the treatment marshes (3), EW influent (4), Allen pond (5), Gearheart pond (6), Hauser pond (7) and Bay discharge (8), were taken weirs marked by the numbers. Map obtained from Google Earth (August 2020, version 9.0). See Supplementary methods for additional descriptions of the ponds and the treatment train.

This region experiences mild seasonal changes throughout the year which are categorized as a wet and dry season. The rainy season averages 100 cm in Winter (December to February), while the dry season (May to September) experiences as low as 12 cm of rain (Table 1, National Weather Service). However, the winter season was unusually dry in 2019, compressing the seasonal periods (TableS1 precipitation). Based on these parameters we consider our study to encompass three seasons: Autumn (October 2019), Winter (November 2019-early January 2020) and Spring (late January 2020). We sampled the water column every 2 weeks to account for these seasonal periods for a total of 8 sampling dates.

**Table 1.**
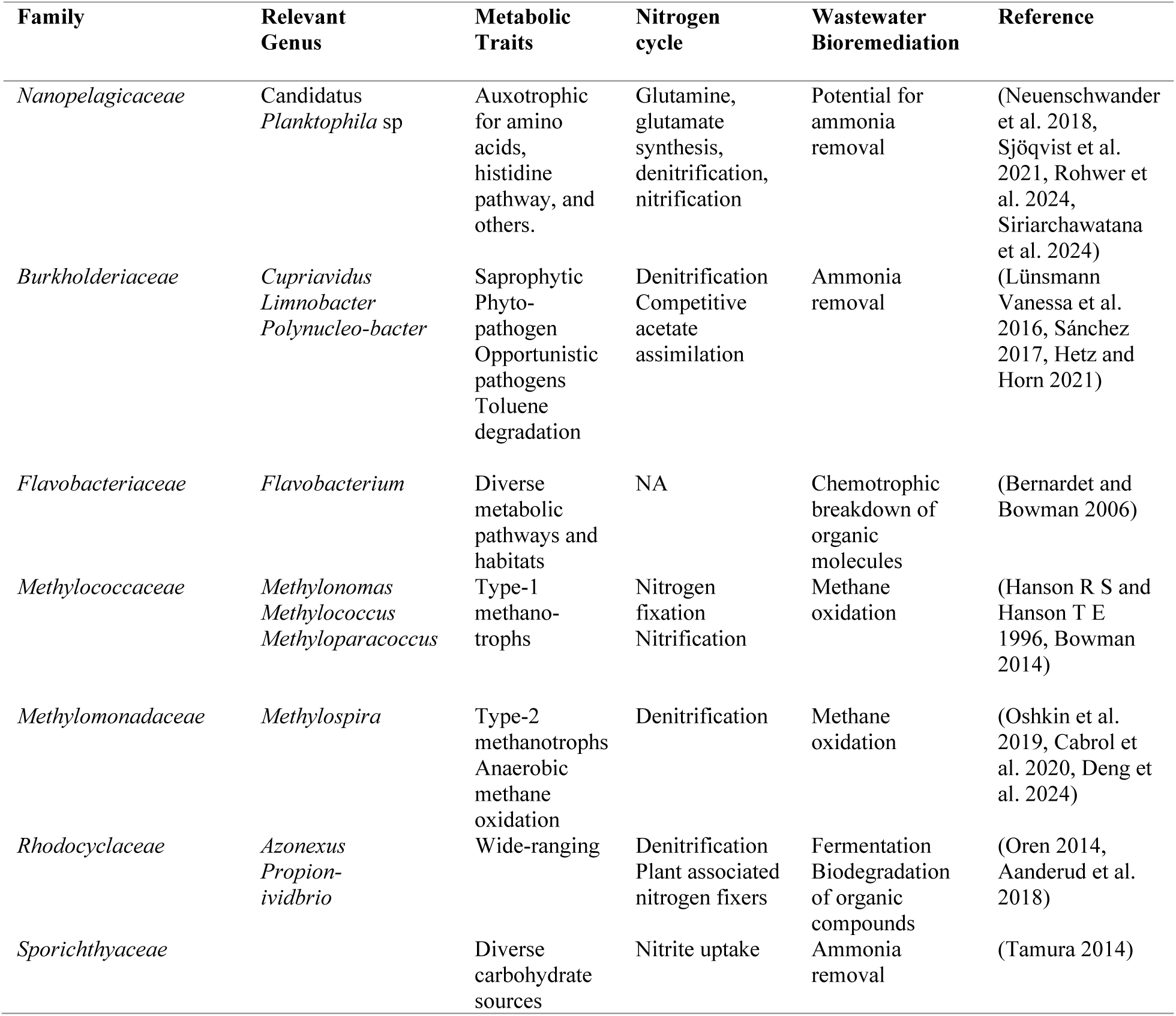
Nitrogen cycle and bioremediation associated traits for the top seven most abundant taxa at the Family level.

Sampling points were chosen to profile how the microbial community and water quality changed as water entered and exited each pond and included all 7 wetland sites in the system (Fig 1). At each sampling date, we took a total of 17 samples that included five samples on each of the two oxidation ponds, two samples from the treatment marshes, two samples from each of the three enhancement marshes, and one bay discharge sample. Samples from the two large oxidation ponds were obtained by boat and encompassed more samples to obtain a reliable profile of its bacterial community and water quality. For the treatment marshes samples, the effluent weir from Oxidation Pond 2 was selected as a representative treatment marsh influent sample, and the effluent was collected as a composite sample at a pre-existing pipe that served as a common effluent for the treatment marshes. The following three enhancement wetlands were sampled via weirs and pump stations chosen to match sampling points used regularly for monitoring conducted by the Arcata Marsh Research Institute (AMRI). For each sampling site, we recorded the water temperature, and dissolved oxygen (DO) using a Hannah multiparameter probe at 3 depths (surface, 30 cm, 90 cm deep) for each location.

Our study focuses on water column bacteria as key representatives for ammonia removal, following the approach used in numerous previous studies (Dong and Reddy 2012, Oopkaup et al. 2016, Sánchez 2017, Morrison et al. 2020). The water column provides a dynamic (Grandmont-Lemire 2022), environmentally sensitive (Zhang et al. 2023) habitat with a diverse heterotrophs community in a relatively homogeneous habitat (Downing and Nerenberg 2007). In this system, the aquatic habitat offers a much larger volume for nitrogen transformation than biofilms and the best point of comparison across heterogeneous ponds.

### Collection and Filtration

Water samples were collected with autoclaved sterile Nalgene 1L bottles, stored in ice and transported to the field laboratory for further processing. For each water sample, 400 *mL* were filtered using sterile analytical filter funnels with a pore size of 0.45 *µm* to minimize clogging from the high algae concentration. Then, the flow-through was collected in a sterile filter flask and filtered a second time through using a pore size of 0.22 *µm* to capture smaller sized bacteria. All processed filters were preserved in 1.5 *mL* microcentrifuge tubes with 1 *mL* of DNase/RNase buffer Zymo Shield (Zymo, Clifornia) and stored at -80° C until ready for extraction.

The remaining sample was used to measured ammonia and nitrate concentration using an Orion meter and the ISE high performance ammonia electrode with substrate specific buffer and 1000 ppm standard solutions (USAbluebook, Illinois). The ammonia test was done with an ammonia probe and calibrated with 100 ppm, 10 ppm, 1 ppm and 0.1 ppm standards.

### DNA Extraction, Sequencing and bioinformatics

We used ZymoBIOMICS DNA/RNA extraction kits (Zymo, California) designed for mixed microbial community samples as described by the manufacturer. The resulting DNA was sequenced using the Illumina MiSeq platform at Argonne National Laboratories. Briefly, the 16S rRNA genes were targeted using archaeal and bacterial primers 515F and 806R, aiming for the V4 region of *Escherichia coli* in accordance with the protocol described in similar previous work (Caporaso et al. 2011, 2012) and applicable by the Earth Microbiome Project (https://earthmicrobiome.org/protocols-and-standards/16s/). The raw DNA sequences have been submitted to the NCBI Sequence Read Archive (SRA) and are accessible under accession number XXXX (provided upon publication).

Raw sequences were demultiplexed using idemp and then quality filtered and clustered using the DADA2 R package (version 1.26, Callahan et al. 2016). Demultiplexed data matching Phi-X reads were removed using the SMALT 0.7.6 akutils phix_filtering command. Chimeras were removed using VSEARCH 1.1.1 (Rognes et al. 2016). Sequences were clustered into Amplicon sequence variants (ASVs) using Greengenes version 13.5 (McDonald et al. 2012) to determine taxonomy. To assign taxa to the ASV table we used the function dada() at the 97% similarity. Using the Phyloseq package (version 3.9, McMurdie and Holmes 2013), we removed ASV assigned to Archaea, Mitochondria, unassigned taxa and those shorter than 1800 number of reads. We used Cumulative Sum Scaling to normalize the resulting reads using the cumNorm() function from the Metagenomeseq package (version 3.19, Paulson et al. 2013).

### Data and Statistical Analysis

All analyses and calculations were completed in R (version 1.4.1564, R core Team 2023) and all graphs and plots were constructed with package ggplot2 (version 2.5.2, Wickham 2016). Custom scripts are available in github (website provided upon publication).

We calculated diversity metrics from the processed ASV table to obtain richness and Pielou’s evenness (Pielou 1959) using functions specnumber() and diversity() in the vegan package (version 2.6-8, Oksanen et al. 2018). We averaged the replicate samplings conducted in Oxidation pond 1 and 2 per date, to account for their large size and maintain equal sampling across ponds. To establish a metric of microbial function, we calculated the delta ammonia concentration of ponds for each sample date by subtracting the effluent ammonia concentration of each location from its influent concentration (Equation 1).

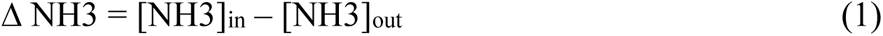

We used a model selection approach to determine the role of richness, season and location in driving Ammonia removal (delta ammonia) as calculated using equation (1). We included a linear model with richness as fixed factor, and generalize linear mixed models that sequentially added season as a fixed factor and location as random factor. This approach accounts for the repeated measurement of each pond, which compromises their independence. We compared linear models using the Aikaike Incormaiton Criterion (AIC) and Bayesian Information Criterion (BIC). Because both criteria indicated the same patterns, we only used AIC in subsequent analyses. We evaluated the best performing model using a Wald test with the function Anova() from the car package in R (Fox and Weisberg 2019). To account for AIC comparisons between linear and GLMM models, we confirmed our findings by comparing linear to GLMM models using analysis of variance with the anova() function in the base R stats package following Galwey ‘s (2014) mock random variable approach.

To reveal diversity and function independent responses to location and season, we used a similar model selection approach as described above, using the Aikaike Information Criteria (AIC) as selecting parameter. We included linear models with season as an explanatory factor and GLMM with season as a fixed factor and location as a random factor. We conducted a Wald test on the best performing model and confirmed our findings by comparing models as described above.

We used Structural Equation Modeling (SEM) to account for the simultaneous influences of location and season on richness and delta ammonia. SEM can be used in ecology to estimate the relationships between multiple dependent and independent variables and better model predictor variables (Quinn and Keoughh 2002, Fan et al. 2016). Our *a priori* model is based on the above GLM and ANOVA approaches, and includes location and season as exogenous variables, while richness and delta ammonia are endogenous variables. We include the direct effect of richness on delta ammonia. We implemented the model using the function sem() from the package lavaan in R (version 06-17, Rosseel 2012). For this analysis, only sites that allowed us to match diversity and function were included (thus excluding OxPond 1 and the influent point), categorical parameters (location, season) were encoded as nominal ordered values to preserve sequence, and numerical parameters (richness, delta ammonia) were scaled and centered at zero. To compare the covariance structure in the model with the covariance matrix of the data we used four goodness of fit methods: χ^2^, root-mean-square error of approximation (RMSEA), CTI and TLI (Iriondo et al. 2003, Tomer and Pugesek 2003).

We assessed the variation of environmental parameters during our study and across the AWTF. The variation of temperature and precipitation are evaluated with linear models since the connectivity between ponds is less significant for these parameters. Season and location were included as factors. Temperature and precipitation were regressed against one another to assess co-variation and, since we found correlations, only the direct effect of temperature was assessed on diversity metrics using linear models. To determine whether the total concentration of ammonia changed with season and location, we used a GLMM with a gaussian distribution that included season as a fixed factor and location as a random factor. We evaluated the significance as indicated above for other GLMMs in this study.

To consider the influence of community composition on BEF relationships, we used a perMANOVA to establish the effects of season and location on microbial composition with the function adonis2() from the package vegan. We further established the independent contribution of season and location using the pairwise.adonis() function (Martinez Arbizu 2020)). For this analysis, we used taxa with 100 or more reads to focus on ASV that were consistently represented in our dataset. We then assessed whether these patterns were driven by differences in community similarity by calculating the Bray-Curtis dissimilarity index and running a multivariate analog to Levine’s homogeneity of variances, using the function betadisper() from the package vegan. We ran a Non-metric Multidimensional Scaling (NMDS) with three dimensions to illustrate these patterns.

We established the taxonomic groups underlying community and functional patterns using a combination of statistical modeling and heatmap imaging. We used a linear model with delta ammonia as dependent variable and ASV relative abundance as explanatory variable to identify potential taxa contributing to function. We focused on the 350 most abundant ASVs, accounting for 75% of the abundance in our dataset and with more than 1600 reads in total. We adjusted p values using the Benjamini & Hochberg (1995) method to account for multiple testing (Cabin and Mitchell 2000). We selected taxa with significant associations with Ammonia removal and performed a GLMM with gaussian distribution, season as a fixed factor and location as a random factor, to establish how these taxa abundances vary across our dataset. We also conducted this modeling at the taxonomic level of family. Significance and corrections were performed as above. To illustrate broader patterns in community composition shifts across location and season, we used the function geom_tile() in the ggplot2 package.

## RESULTS

After quality filtering, we obtained 3 945 058 reads distributed across 137 samples with an average of 29 007 reads per sample (+-6864), and clustered in 8 702 ASVs from all wastewater samples combined throughout the treatment plant for the duration of the study. The taxa belonged to 47 Phyla, 438 Families and 753 Genera from the Kingdom Bacteria, with only 9 attributed to Archaea. In the following paragraphs, we will show the broader BEF relationship, the effect of location and season on ASV diversity and ammonia removal function, the environmental conditions potentially underlying the effect of location and season, and the effect of community composition, while identifying taxa associated with ammonia removal.

### Overall BEF relationship

We found a significant negative BEF relationship between richness and function when accounting for all samples across the wastewater treatment plan for the duration of the study (Fig 2). The best performing model included only richness as an explanatory factor (F_1_ = 5.251, p=0.030, Fig 2) and outperformed any models that included season or location (Table S2a). Samples at the lowest range of the richness gradient had in average 101.5 ASVs and corresponded to the removal of 8.305 mg/L Ammonia. In contrast, samples at the highest range of the richness gradient had, in average, 940.68 ASVs, but only accounted for 5.315 mg/L Ammonia removal. There was no significant relationship between delta ammonia and species evenness (GLMM: χ ^2^ = 0.760, *df* = 1, *p*-value = 0.383). The best performing model to explain function including evenness as a fixed factor included location as a random factor but season was not included (Table S2b).

**Figure 2.**
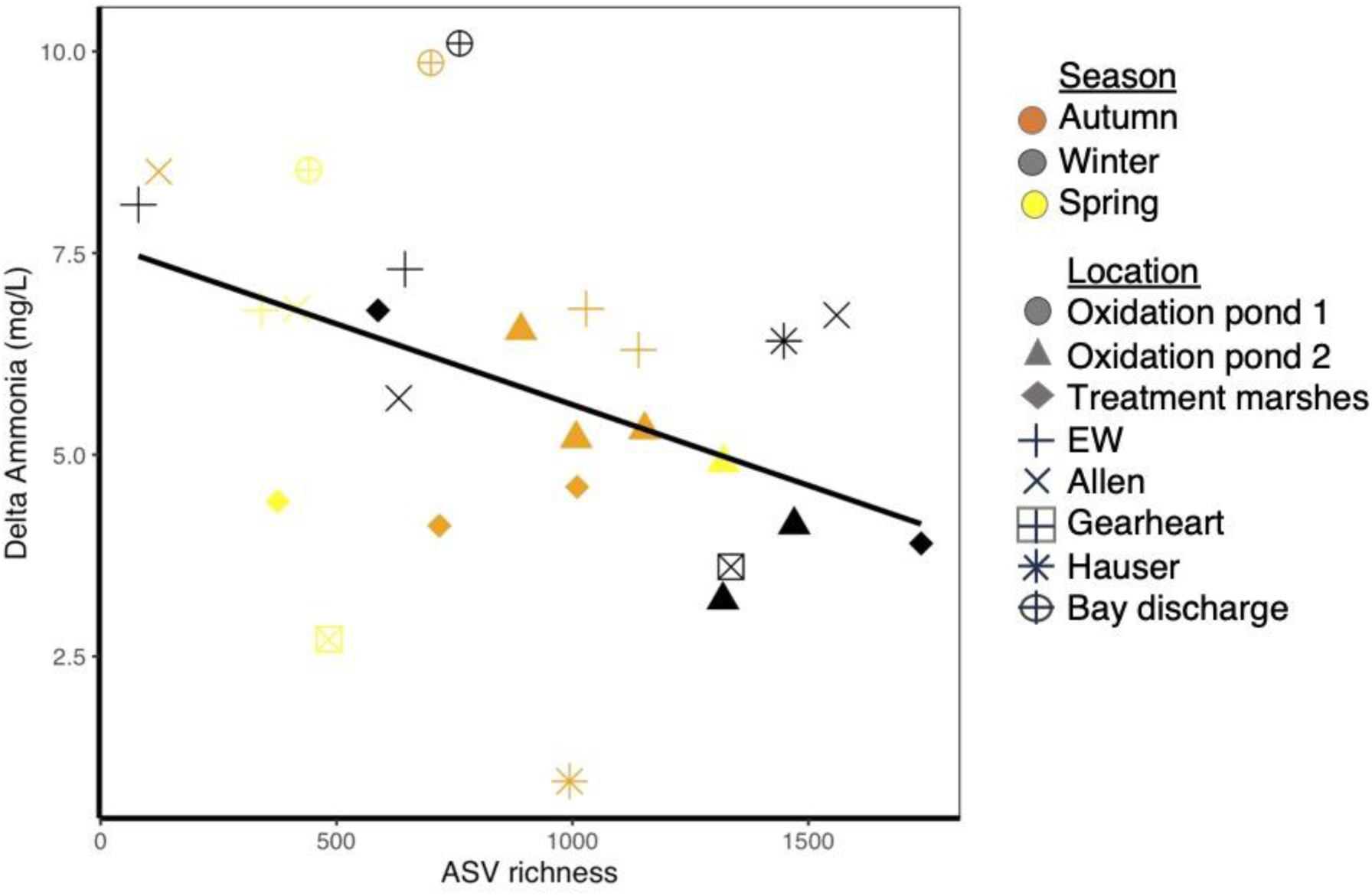
Negative BEF relationship between diversity and the average change in ammonia for each pond at each date. The line represents the linear relationship between Richness and Delta Ammonia.

### Effects of location and season

The function of ammonia removal (Delta Ammonia) responded to location but not to seasonal change (Fig 3). The best performing model included only location as a random factor to explain Delta ammonia data (Table S3). We tested the GLMM with only random factors using a mock random fixed variable to be able to use the Wald test. Although the model was not significant (χ^2^=1.358, df=1, p=0.244), it performed better than the null model (χ^2^=24.972, df=1, p<0.001). The lack of significance may be related to the pattern of increase and decrease in Ammonia removal (mg/L) along the series of treatment marshes, with higher values in Allen Pond and at the bay discharge point (Fig 3). Season was not included in any of the best performing models (Table S3).

**Figure 3.**
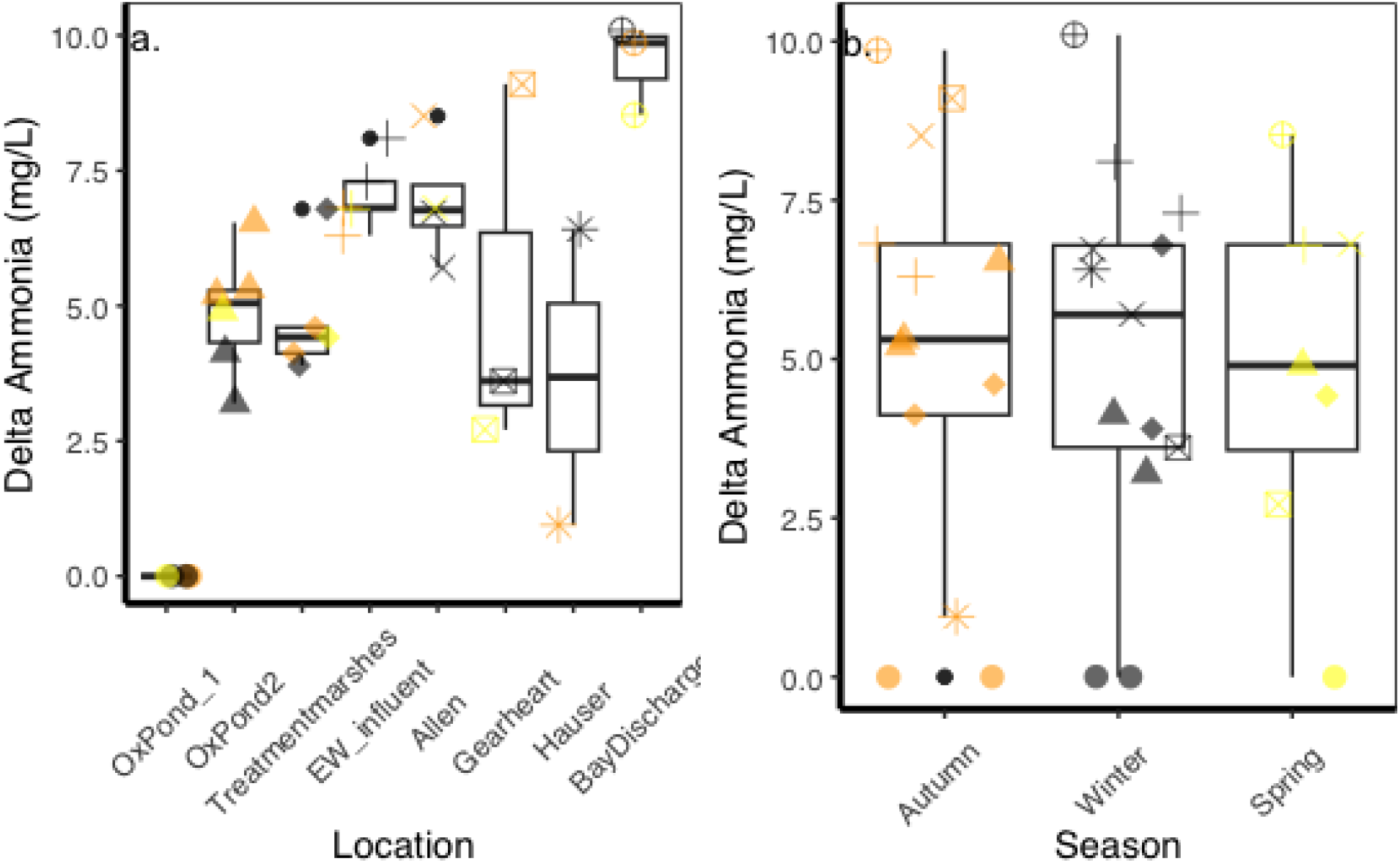
Effects of location (a) and season (b) on the change in ammonia concentration. In (b), locations are ordered from the input to the bay discharge point. See figure X for legend details.

ASV richness responded to season and, only weakly, to location. Season had a significant effect on richness (χ ^2^=8.1269, df=2, p=0.0172). Winter had the highest ASV numbers (1256 taxa), declining towards Spring (603 taxa, Fig 4). The contribution of location was not significant, even though the best performing model included the random variable (Table S4a). Notably, Oxidation pond 2 and Hauser had the highest mean richness, while the lowest averages were found in Gearheart and Bay discharge points. These low points corresponded with the chlorination steps in the treatment train. Allen marsh had the highest variability in richness but was not significantly different from the other locations (Fig. 4). Evenness showed a similar pattern with Season as the main contributing factor, but the best performing model was a linear model with only Season as a fixed factor (Fig S1, Table S4b, Wald test: F_2_=6.1714, p=0.004).

**Figure 4.**
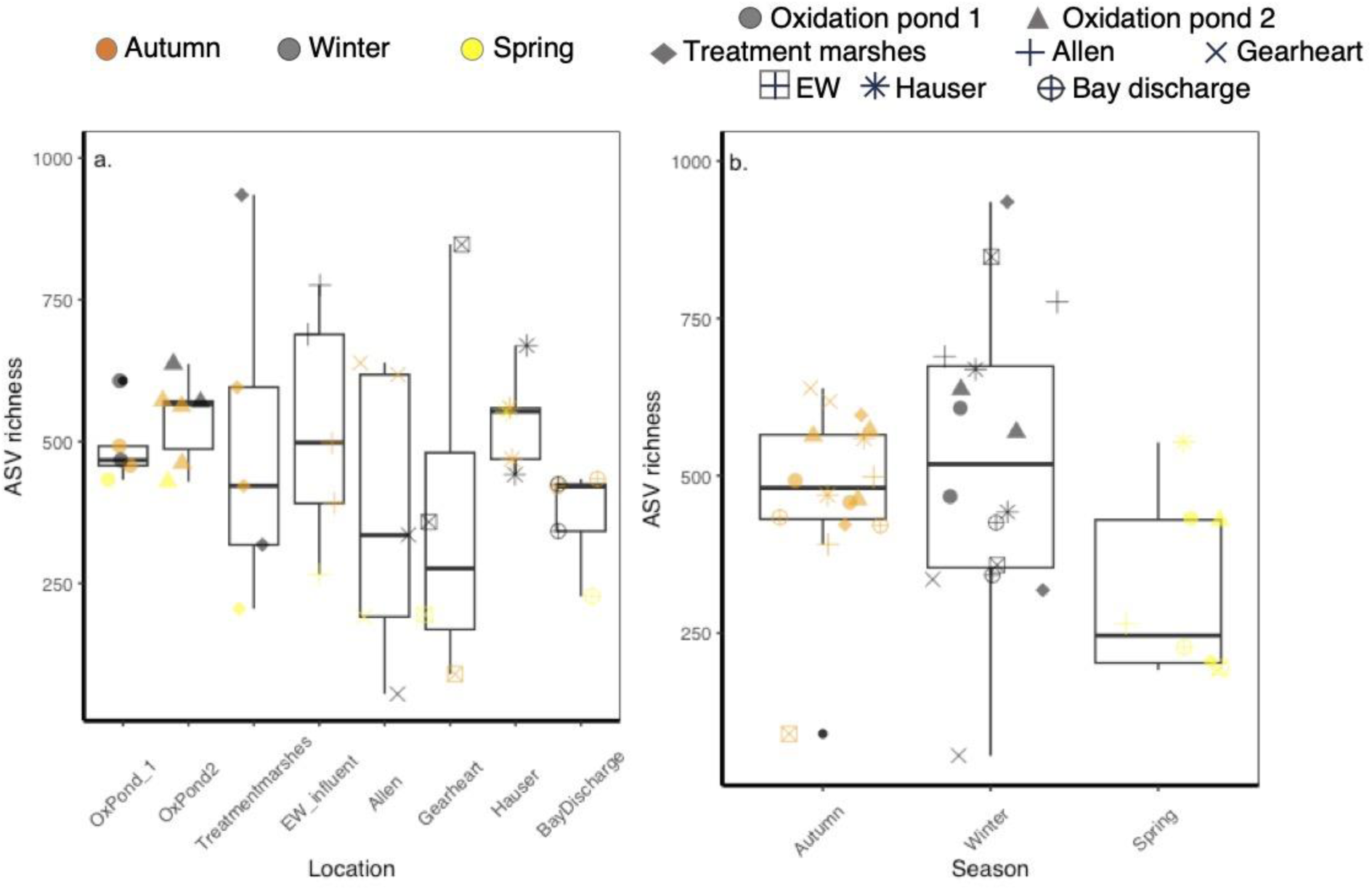
ASV richness responses to a) location and b) season. Datapoints represent individual samples at each sampling event. Colors represent Seasons in (a) while shapes represent locations in (b). Oxidation pond data was averaged to account for their large sizes and multiple sampling.

To better understand the influence of space and season on bacterial diversity and delta ammonia, while taking into account the effect of diversity on function, we used a Structural Equation Model (SEM). As with the GLM models, we found that richness had a negative effect on Delta ammonia (coefficient = -0.28). The SEM also revealed negative effects of season (coefficient = -0.08) and location (weaker, coefficient = -0.16) on richness (Fig 5). In contrast, Delta ammonia was influenced negatively by season, but positively by location (coefficient= 0.21). The magnitude of some of these coefficients seems at odds with the results we obtained from the GLM and ANOVA analyses, likely due to the locations we had to exclude from the SEM analysis. The selected goodness of fit assessment tools indicated strong support for the model (CFI=1.00, TLI=1.00, RMSEA=0) except for Chi2 (Chi2 =0, p=0), which is known to be highly sensitive to sample sizes (Fan et al. 2016).

**Figure 5.**
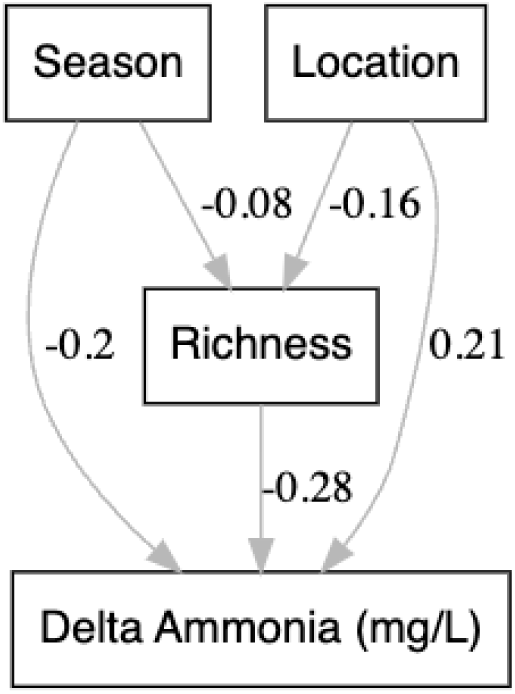
Structural equation model (SEM) showing combined effect of season and location on diversity and function. Arrows represent relationships between measured variables (rectangles). Coefficients are indicated for each relationship.

### Abiotic parameters associated with location and season

Temperature and precipitation seemed to be representatives of seasonal change while total Ammonia levels was closely related to sites along the series of treatment marshes. Temperature declined from Autumn to Winter, but did not change significantly with location (Fig S2 top panels, R^2^=0.839, F_22, 10_ = 8.589, p < 0.001). Precipitation followed a similar trend, increasing from Autumn to Winter, and showing no response to location (Fig S2 middle panels, R^2^=0.914, F_22, 10_ = 14.29, p < 0.001). Because temperature and precipitation are weakly correlated (R^2^=0.78, F_1,31_=117.4, p<0.001), we only test richness against one of the parameters, temperature, and find no significant relationship (Fig S2 bottom panels, R^2^=0.05, F_1,31_=117.4, p=0.687). Evenness followed the same trend and was unrelated to temperature (R^2^=0.05, F_1,31_=2.772, p=0.106).

Total levels of ammonia declined throughout the system (Fig S3) from Oxidation Pond 1 (25.08 mg/L) to the Bay Discharge (2.79 mg/L) with a slight peak at the EW influent (26.87 mg/L, fig ammonia). Overall, ammonia concentration ranged from 40 mg/L at Oxidation pond 1 in the winter to 2.79 mg/L at the Bay discharge in the spring. The best performing model included season as a fixed factor and location as a random factor (Table S5). Although the model was not significant after a Wald test (χ ^2^=1.354, df=2, p=0.508), it was significantly better than the null model (χ ^2^=25.055, df=2, p<0.01) and performed better than the linear model with season as fixed factor (Table S5). Unfortunately, pH, BOD and DO measurements were compromised due to field conditions and technical difficulties, thus they were not included in any further analysis.

### Bacterial composition and representative taxa

Bacterial community composition responded to location and season (Fig 6, perMANOVA: R^2^=0.306, F_23,89_ = 1.711, p = 0.001). The effect of Season was maintained after a pairwise test (F_2,89_ = 8.133, p=0.02), but the effect of location was lost (F_7,105_ = 1.398, p=0.098). Bacterial communities seemed more similar during Spring than other seasons even though we did not find significant differences (betadisper: F_2_=2.877, p=0.060). Location had no effect on community similarity (F_7_=1.099, p=0.369).

**Fig 6.**
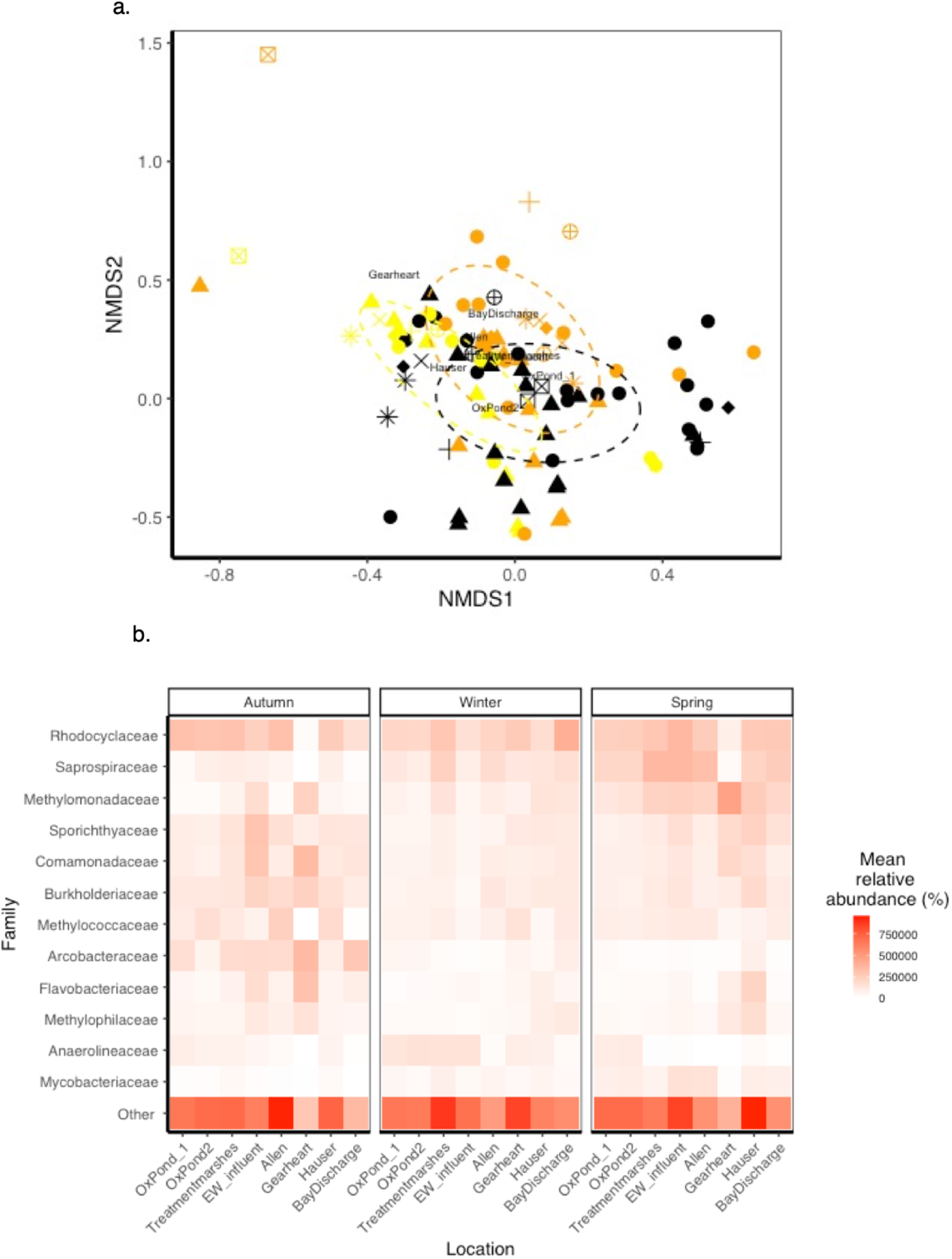
Community composition responses to location and season. (a) NMDS plot of ASV composition by season (color) and location (shape). The best solution for the NMDS had a stress of 0.1458 with a Procrustes RMSE of 0.010 and a maximum residual of 0.095. See Fig 2 for legend details. Lines depict the covariance ellipses for season. Centroids for location are indicated with the name of each pond. (b) Heatmap of average relative abundances at family level taxonomic distribution. Shown are the most 13 abundant families in the dataset with other taxa clustered in ‘others’. See Fig S4 for an extended version of this figure. Darker tones indicate higher relative abundance.

Out of the original 346 ASVs, we found 11 taxa with positive influence on Delta Ammonia, 331 taxa with neutral or no association and 3 taxa with negative association (Table S6). Taxa with positive associations with ammonia removal included those classified as *Legionella sp* and Candidatus *Planktophila sp,*(Arcobacter) and belonging to Sulfurimonadaceae, Sporichthyaceae, Rhizobiales, Rhodocyclaceae, and Oxalobacteraceae. Taxa with negative association with function included members of Omnitrophaceae, Methanosaetaceae and Cloacimonadacea. Most of these taxa’s abundances changed with location and season, except for Cloacimonadaceae, which only responded to season (Table S7). The qualitative analysis at the family level also revealed an interaction between location and season (Fig 6b). Of the taxa identified above, only Rhodocyclaceae, Sporichthyaceae and Arcobacteriaceae were numerically dominant (Fig 6b). At the family level, three families were identified to have significant correlation with ammonia removal: Methylococcaceae, Burkholderiaceae and Legionellaceae (Table S8). Some families increased in abundance with season, including Mycobacteriaceae, Methylomonadaceae, and Saprospiraceae. A few groups decreased in abundance as seasons changed including Rhodocyclaceae, Sporichthyaceae, Comamonadaceae, and Burkholderiaceae.

Other groups varied from site to site and across seasons, making inferences challenging. For example, Arcobacter generally declined with season but it was very variable across sites in Autumn. Methylocaoccaceae was abundant in Oxidation pond 2, Allen and Hauser marshes, but only in autumn samples, with a significant association with ammonia removal (Table S6). Notice that rare taxa are very abundant and vary across sites and seasons, highlighting the role of rare species.

Notably, Allen and Gearheart Ponds had the highest variability in family and genera composition, with a striking response to season. These ponds are lacking taxa that are otherwise abundant in the system, such as Rhodocyclaceae and Saprospiraceae in Gearheart Pond, and Methylomonadaceae and Flavobacteriaceae in Allen Pond, especially in Autumn and Spring (Fig 6b). In contrast, these sites are enriched with taxa such as Methylococcaceae, Bulkholderiaceae, Comamonadaceae, Flavobacteriaceae and Arcobacteraceae, particularly during autumn.

## DISCUSSION

A key goal in ecology today is to manage and sustain resilient ecosystem functions that persist in a human-influenced changing world (Balvanera et al. 2006, Bodelier 2011, Hautier et al. 2014). Steps forward should include a dynamic view of Biodiversity-Ecosystem function relationships that integrates abiotic and biotic processes shaping community diversity, composition, and functional outputs across space and time. We tested this framework in a constructed wetland used for wastewater secondary treatment and found a negative relationship between ammonia removal and bacterial diversity (Fig 2). This pattern emerged from seasonal effects on bacterial diversity and location-specific influences on function (Fig 3), highlighting the importance of considering the interaction between environmental, spatial and biological factors. Our work thus adds to the growing body of research emphasizing the ecological context of BEF relationships (Grace et al. 2016, Berlinches de Gea et al. 2023) which often do not account for spatial connectivity or report negative relationships.

Negative biodiversity-ecosystem relationships are rare (van der Plas 2019, D’Andrea et al. 2024) and have primarily been documented for microbial degradation within aquatic (Jiang 2007, Peter et al. 2011, Cuellar-Gempeler et al. in review) and soil systems (Toljander et al. 2006). These negative relationships are explained in the literature by the negative selection effect, which suggests that species contributing to function are poor competitors, or subordinate (Jiang 2007, Jiang et al. 2008). Our findings suggest that bacteria with the highest potential for ammonia removal are subordinate species thriving in low-diversity locations after chlorination steps in the treatment train, supporting Hypothesis 1. The strong influence of chlorination on microbial communities (Murray G E et al. 1984), seems to be a disturbance point that overrides other aspects of connectivity and heterogeneity in this system. We propose that these disturbances create a refuge for bacterial taxa that support the highest ammonia removal rates in the system.

These disturbance points of ammonia removal refugia are clearly illustrated by the seasonal patterns in Allen and Gearheart ponds. These two sites have the lowest richness values (Fig 3) and are lacking in taxa that are abundant otherwise in the system. They instead host taxa are involved in critical steps within nitrogen cycling beyond just denitrification (Table 2), including Bulkholderiaceae, Rhodocyclaceae, and Flavobacteriaceae. These patterns underscore the critical role of specific locations in facilitating ammonia removal from the system as functional refugia, which, to our knowledge, not been reported in the context of wastewater treatment marshes. Notably, the effects of chlorination on Allen and Gearheart ponds are season-dependent, probably because the chlorination effect weakens during Humboldt’s wet winters brings higher flow rates into the AWTF, which dilute the impact of chlorination.

Seasonal change resulted in increased diversity during the wet winter months (Fig 3) and had no significant effects on function (Fig 2), thus refuting our Hypothesis 2. Instead, functional redundancy may explain this pattern, as taxa with similar functional roles replace one another as seasons shift, maintaining ammonia removal despite taxonomic turnover (Walker 1992, Lawton 1994, Loreau 2004). For example, Burkholderiaceae, known competitive acetate assimilators during complete denitrification (Hetz and Horn 2021), decreased slightly from Autumn to Spring (Table S7) while several members of Methylococcaceae became abundant in Spring and can play key roles in denitrification and methane flux, known as methanotrophic denitrification (Dedysh et al. 1998, Reis et al. 2024). Additionally, macrophytes and algae can buffer seasonal change for bacteria by providing substrate and oxygen during winter months, and carbon sources during summer months. We find a general negative BEF relationship with evidence of seasonal redundancy, suggesting redundancy can be context-dependent, perhaps contributing to an ongoing controversy regarding redundancy in microbial ecology. Traditionally, it was assumed that high diversity in microbial organisms implied functional redundancy (Allison and Martiny 2008), yet recent empirical evidence support positive BEF relationships suggesting complementarity (Delgado-Baquerizo et al. 2016, Galand et al. n.d.). Perhaps we should shift focus from one general BEF relationship and instead establish when to expect different BEF relationships based on ecological contexts and biological parameters.

Our SEM analysis revealed that bacterial richness had stronger influence on ammonia removal than season or location (Fig 5), underscoring the biological basis of this function as primary mediator of environmental effects. The direct influence of location on ammonia removal suggests that site-specific conditions may impact per capita functioning or regulatory mechanisms, independent of bacterial population dynamics. Some of these site specific-effects include the impact of chlorination, the contribution of macrophytes to bacterial activity, and the declining ammonia levels across the treatment train. The role of per capita functioning in shaping ecosystem processes is understudied (Leander et al 2016) and may be crucial to understand how environmental factors influence ecosystem functions like primary productivity (Chalifour and Juneau 2011) and nitrogen cycling (Vitousek et al. 1997). Our model also suggests season may have per-capita effects on function, while their direct impact on richness -linked to population shifts-appears weaker (Fig 5). This combination of direct and indirect effects highlights a close and intricate interplay between environmental drivers, diversity and functional output in wetland ecosystems. Our results align with previous work that integrates environmental context, site disturbances and heterogeneity (but not connectivity) and species diversity into models that highlight the multifaceted drivers of ecosystem function (Grace et al. 2016, Martins et al. 2024). In contrast, some studies have shown stronger environmental effects on microbial functionality than biodiversity (Jing et al. 2015, Delgado-Baquerizo et al. 2016, 2020) likely due to their focus on multifunctionality (Byrnes et al. 2014).

Although beyond the scope of this study, there are two factors that can influence bacterial diversity and function in constructed wetlands that we did not address. First, Dissimilatory Nitrogen Reduction to Ammonia (DNRA) and nitrogen fixation are metabolic pathways that can maintain ammonia in marsh systems contributing to overall ammonia budget in wastewater treatment (Pandey et al. 2020). Our system likely does not support important DNRA activity due to its declining organic load (following Oxidation pond 1) and low sulfide levels. While nitrogen fixation may occur, we did not detect large abundances of typical nitrogen fixing taxa in in our dataset. Second, studies often emphasize the role of plant-microbe interactions and biofilms in wetland water purification (Brisson et al. 2020). Wetland rhizosphere creates oxic-anoxic interfaces that stimulate nutrient cycling (Vacca et al. 2005, Gagnon et al. 2007). We acknowledge that plant and wildlife heterogeneity between ponds may influence bacterial diversity and function but recognize the technical challenges in measuring these variables for the purpose of this study. Sediment and detritus may contribute to increase ammonia in the system via the slow process of degradation and decay. We instead focused on the water fraction as a way to facilitate comparisons across with varying vegetation, given that volume of the aquatic habitat is larger and more impactful in our system.

Overall, our findings coincide with previously reported ammonia levels, ammonia removal rates, bacterial diversity and composition. We found ammonia levels that ranged from 2.79 to 40 mg/L and declined consistently as the water moved from the influx at Oxidation pond 1 to the bay discharge point (Figure S3). These results are comparable to constructed wetlands for domestic wastewater treatment reported before (Lu et al. 2014, He et al. 2016). The community structure and dominant taxa we found were similar to previously reported work and included Rhodocyclaceae (Zheng et al. 2022), Flavobacteriaceae (Adrados et al. 2014, Sidrach-Cardona et al. 2015, Zhong et al. 2015, Guan et al. 2015, Xu et al. 2016), Saprospiraceae (Ginige Maneesha P. et al. 2004, Sun et al. 2012, Zhang et al. 2023), Methylococcaceae (Siniscalchi et al. 2017) and many others (Sánchez 2017) with critical functional roles (Table 2). Interestingly, one of the taxa with strongest associations with ammonia removal, Candidatus *Planktophila* sp. (Actinobacteria, Table 2) is an unculturable auxotroph, hypothesized to participate actively in nitrogen cycling despite its small genome (Sjöqvist et al. 2021, Rohwer et al. 2024, Siriarchawatana et al. 2024). *Planktophila* sp. has been only recovered when grown in mixture because it requires aminoacids and carbon metabolism intermediates (Neuenschwander et al. 2018). Procurement of such growth factors may be accomplished by *Legionella* sp in our study, a taxon that was positively associated with function but with no evidence of nitrogen cycling functions. This highlights the role of species contributing to function and their critical interactions with other members of the community.

In contrast to the similarities outlined above, our findings deviate from some studies due to critical differences in hydraulic design (Kadlec 2009, Vymazal 2011, Liao et al. 2013, Sidrach-Cardona et al. 2015, He et al. 2016), due to more extreme seasonal patterns (Garfí et al. 2012, Sánchez 2017), focus on functions other than ammonia removal, and complex industrial or polluted influx (Abed et al. 2014, Guan et al. 2015). It seems like the unique environmental, operational and biological conditions of each wastewater treatment wetland results in case-specific management requirements and challenges (Sánchez 2017).

Despite this individuality, three general recommendations stemming from our findings include to (1) consider BEF theory when establishing relationships between biodiversity and purification functions, (2) identify functional refugia and focus monitoring goals on those conditions, (3) recognize functional redundancy and its buffering potential for climatic fluctuations. Wastewater treatment constructed wetlands represent a unique opportunity to apply and test our understanding of ecological processes and use microbial activity to reduce anthropogenic environmental impact. We propose that the concepts of functional refugia and seasonal functional redundancy may be useful in the context of a dynamic BEF field that recognizes the role of environmental context.

## Acknowledgements

We gratefully acknowledge the funding provided by the Arcata Marsh Research Institute, Dr. Cuellar-Gempeler’s Microbial Laboratory, the City of Arcata, and scholarships from the Humboldt Marine and Coastal Science Institute, which made this project possible and facilitated the use of advanced molecular techniques. We also extend our thanks to the project’s committee members for their valuable insights and expertise. Additionally, we would like to express our appreciation to the members of the lab for their support, collaboration, and contributions throughout the project. Their dedication, assistance with experimental procedures, and thoughtful discussions were essential to the success of this work.

